# Mcam stabilizes luminal progenitor breast cancer phenotypes via Ck2 control and Src/Akt/Stat3 attenuation

**DOI:** 10.1101/2023.05.10.540211

**Authors:** Ozlen Balcioglu, Brooke L. Gates, David W. Freeman, Berhane M. Hagos, Elnaz Mirzaei Mehrabad, David Ayala-Talavera, Benjamin T. Spike

## Abstract

Breast cancers are categorized into subtypes with distinctive therapeutic vulnerabilities and prognoses based on their expression of clinically targetable receptors and gene expression patterns mimicking different cell types of the normal gland. Here, we tested the role of Mcam in breast cancer cell state control and tumorigenicity in a luminal progenitor-like murine tumor cell line (Py230) that exhibits lineage and tumor subtype plasticity. Mcam knockdown Py230 cells show augmented Stat3 and Pi3K/Akt activation associated with a lineage state switch away from a hormone-sensing/luminal progenitor state toward alveolar and basal cell related phenotypes that were refractory to growth inhibition by the anti-estrogen therapeutic, tamoxifen. Inhibition of Stat3, or the upstream activator Ck2, reversed these cell state changes. Mcam binds Ck2 and acts as a regulator of Ck2 substrate utilization across multiple mammary tumor cell lines. In Py230 cells this activity manifests as increased mesenchymal morphology, migration, and Src/Fak/Mapk/Paxillin adhesion complex signaling *in vitro*, in contrast to Mcam’s reported roles in promoting mesenchymal phenotypes. *In vivo*, Mcam knockdown reduced tumor growth and take rate and inhibited cell state transition to Sox10+/neural crest like cells previously been associated with tumor aggressiveness. This contrasts with human luminal breast cancers where MCAM copy number loss is highly coupled to Cyclin D amplification, increased proliferation, and the more aggressive Luminal B subtype. Together these data indicate a critical role for Mcam and its regulation of Ck2 in control of breast cancer cell state plasticity with implications for progression, evasion of targeted therapies and combination therapy design.

**HIGHLIGHTS:**

- Mcam is expressed by diverse breast cancer subtypes.
- Mcam regulates breast cancer cell state plasticity.
- Mcam maintains the mammary luminal progenitor state.
- Mcam attenuates Stat activity.
- Mcam dictates Ck2 substrate utilization.

## 1. INTRODUCTION

Breast cancers exhibit cellular and subtype heterogeneity that complicate treatment and contribute to disease progression. Whereas hormone receptor positive (HR+) and Her2 amplified (Her2+) cancers are treated with molecular therapies targeting their molecular drivers, triple negative breast cancers (TNBC) lack these molecular targets.^3^ Transcriptional profiling has identified intrinsic breast cancer subtypes that partially overlap with these clinical subtype designations and with cellular compartments of the normal breast.^4^ For instance, Luminal A and B subtypes are typically HR+ (estrogen and/or progesterone receptor expressing), the ‘Her2-enriched’ intrinsic subtype comprises most Her2 amplified tumors, and the ‘Basal-like’ designation encompasses most TNBCs.^5^

The origins of this heterogeneity remain controversial, and the molecular mediators are incompletely understood. However, it is generally held that tumor subtype reflects particular combinations of driver mutations and intrinsic characteristics of the cell of origin.^3,6,7^ As such, experiments in mice targeting drivers to different mammary gland cell types give rise to different subtypes of breast cancer.^8,9^ Nevertheless, cell plasticity has emerged as a significant contributor to intra- and inter-tumoral heterogeneity, likely playing a crucial role in therapy resistance, dormancy, and metastasis in breast and other cancers.^10,11^ Examples include epithelial-to-mesenchymal transition (EMT)—a classic plasticity mechanism implicated in tumor cell migration, invasion, and metastasis^9,12^—and cell lineage switching within the epithelial compartment, a well-documented mechanism of therapy evasion and progression in diverse cancers.^13–15^ Thus, hormone-dependent cancers can sometimes transition into hormone-independent phenotypes that evade conventional hormone-blocking therapy. In breast cancer, this conversion has been associated with EMT or alternatively a state transition to a hormone insensitive, alveolar progenitor program classically driven by Elf5, and Stat3/5.^16–18^ Identifying and understanding the molecular regulators of epithelial lineage plasticity in breast and other cancers could reveal strategies to maintain therapy sensitivity or uncover new therapeutic vulnerabilities. In this regard, we chose to examine the membrane-integral, cell-surface glycoprotein MCAM (CD146; MUC18; S-Endo-1; and Gicerin) as a regulator of epithelial plasticity in breast cancer based on several prior observations. Our prior studies showed that Mcam is broadly expressed in multipotent fetal mammary stem cells.^1^ Mcam is an established marker of pericytes noted for their high intrinsic cellular plasticity, although lower-level expression in a variety of other cells and tissues, including multipotent mammary stem cell and luminal progenitor (LP) compartments, has been reported.^19,20^ MCAM is upregulated in a variety of cancers and has different activities depending on the target cancer/cell type.^19,21^ In breast cancers, elevated expression of MCAM has been associated with the Basal-like subtype and poor prognosis.^19,22^ Experimental overexpression (OE) of MCAM in MCF7 and SKBR-3 cells (HR+ and HER2+ cell lines, respectively) drives EMT and enhances tumorigenesis in xenograft settings.^23^ However, MCAM has also been reported to be tumor suppressive and recent work has challenged the notion that it is widely oncogenic.^24^

Mechanistically, MCAM has been implicated in a variety of biological processes relevant to mammary development, cell identity, and cancer. MCAM can mediate cell-cell interactions between heterotypic cells during extravasation of immune and tumor cells and promotes homotypic cell interactions through cognate binding partners that are yet to be identified.^19^ MCAM binds S100A8/9, Galectin-1/3 and Laminin-α4 as extracellular ligands and regulates Integrin β1 and CD44 levels on the cell surface.^25–27^ Inside the cell, MCAM regulates integrin-mediated signaling through PIP2 and FAK activation, and ERM binding, and cooperates in trafficking of a discrete internalized organellar structure associated with Wnt-mediated planar cell polarity.^28–30^ The MCAM cytoplasmic tail possesses putative binding/phosphorylation sites for Src family kinases, PKA, PKC and CK2 that may orchestrate coupling and sequestering of broad-spectrum kinases to and from their targets.^31^ To date, it is unknown how these signals are integrated into cell fate choices, particularly in luminal breast cancer cells where MCAM has thus far only been studied under conditions of acquired or experiment OE.^19,23,29,31^

In the present study, using a LP/stem cell-like mammary tumor cell line derived from the MMTV-PyMT mouse model (Py230) that exhibits lineage and tumor subtype plasticity and comparing this to a number of additional cell lines and human archival tumor data, we found that Mcam is a potent regulator of breast cancer cell state, therapy resistance and tumorigenicity.^9^ Mcam’s effects are mediated by its governance of the master regulatory kinase Ck2. Mcam KD altered Ck2 substrate utilization, including its activation of Stat3 and inactivation of PTEN. Inhibition of Ck2/Stat3 reversed these signaling and transcriptional state changes, and the associated insensitivity to tamoxifen—a common characteristic of human Luminal B breast cancers which we note often harbor MCAM loss. These results may inform the development of combination breast cancer therapies that incorporate MCAM and/or CK2 and STAT inhibitors with existing molecularly targeted therapies and point to opportunities as well as potential challenges in clinical management of breast cancer.

## 2. METHODS

### 2.1. Cell lines and Cell culture

Met-1 and NDL-1 cell lines were provided by Dr. Alexander Borowsky.^32,33^ Authenticated 4T1 (RRID: CVCL_0125), E0771 (RRID: CVCL_GR23), MCF10A (RRID: CVCL_0598), MDA-MB-231 (RRID: CVCL_0062) and MDA-MB-468 (RRID: CVCL_0419) cell lines were obtained from ATCC. 4T1, E0771, NDL-1, Met-1, MDA-MB-231, and MDA-MB-468 cells were cultured in Dulbecco’s modified Eagle’s media (DMEM) with ciprofloxacin 10 μg/ml and 10% fetal calf serum. MCF10A cells were cultured in Dulbecco’s modified Eagle’s medium/F12 with 5% horse serum, 10 μg/ml insulin, 20 ng/ml epidermal growth factor, 100 ng/ml cholera toxin, 0.5 μg/ml hydrocortisone, and 10 μg/ml ciprofloxacin. Py230 cell line was kindly provided by L. Ellies ^9^. Identity of gifted lines is authenticated herein through molecular assays (e.g., scRNASeq,) that match profiles reported for these lines. Passage numbers were tracked and minimized, and experiments were repeated 2-8 times including repetition with early passage frozen aliquots. Py230 cells were cultured in “Py230 Media” composed of F12-Kaighn’s Modified Media with 5% fetal calf serum, ciprofloxacin 10μg/ml, amphotericin B 2.5μg/ml, and Mito+ serum extender (Corning #355006). Py230 organoids were cultured in Py230 media with 4% Matrigel. HCI-011 organoids were kindly provided by Dr. Alana Welm and cultured according to their published protocol.^34^ They were grown in PDxO base medium (Advanced DMEM/F12 with 5% FBS, 10 mM HEPES, 1× Glutamax, 1 μg ml^−1^ hydrocortisone, 50 μg ml^−1^ gentamicin and 10 ng ml^−1^ hEGF) with 10μM Y-27632, 100 ng ml^−1^ FGF2 and 1 mM NAC. All cells were grown in sterile humidified tissue culture incubators at 37 °C with 5% CO_2_ and ambient (∼ 17–18%) O_2_. Cells were transduced with lentiviral concentrates in the presence of 7 μg/ml polybrene for up to 16 h with transduction efficiency (not shown) suggesting MOI <<1. CX-4945/Silmitasertib (Selleck Chem #S2248), (E/Z)-GO289 (MedChem Express, #HY-115519), Ag490 (Selleck Chem #S1143), Stattic/S7947 (S7947, Sigma Aldrich #19983-44-9), and tamoxifen citrate (Selleck Chem #S1972) were resuspended in DMSO according to manufacturers’ recommendations and diluted to final concentrations in media immediately prior to being added to cells.

### 2.2. Lentiviral vectors

Viral vector plasmids were obtained from OriGene Technologies, Inc. (Rockville, MD), including the mouse Mcam shRNA lentiviral plasmid (Cat # TL514377), a scrambled shControl, and a custom shRNA-resistant (via silent-mutation) mouse Mcam construct subcloned into a lentiviral gene expression vector (pLenti-C-mCFP-P2A-BSD), (# PS100107). Completed vectors were sequenced confirmed.

### 2.3. siRNA construct

siRNA constructs were obtained from OriGene Technologies, Inc. (Rockville, MD). This includes the three unique 27mer mouse Mcam siRNA duplexes (SR418656A-C) and a Trilencer-27 Universal Scrambled Negative Control siRNA Duplex (SR30005). Cells were transfected utilizing Lipofectamine 3000 Transfection Reagent (Invitrogen, #L3000008) according to manufacturer’s instructions.

### 2.4. Western Blot

Cell were lysed in RIPA buffer (1X PBS (137mM NaCl; 2.7mM KCl; 4.3mM Na2PO4; 1.47 mM KH2PO4) with 1% Nonidet P-40 Substitute (Sigma #74385), 0.5% Sodium deoxycholate, 0.1% SDS) with the Halt Protease & Phosphatase inhibitor cocktail (Thermo Fisher Scientific, #78440), quantified with the RC-DC Protein Quantification Assay (BioRad) and equivalent concentrations (10-16ug) loaded per sample on precast NuPAGE 4-12% Bis-Tris gradient gels (Invitrogen). Separated proteins were transferred to nitrocellulose and probed overnight with primary antibodies under agitation. Secondary antibodies were incubated with blots for 1hr. Intervening washes used TBST pH7.5 (10mM Tris, 15mM NaCl, 0.05%Tween 20), Blots were imaged on an Odyssey CLx Imager (LI-COR).

### 2.5. Immunoprecipitation

HEK293A cells were transfected with ps14-Mock ^35^, an rTA expression construct to later induce CK2 expression, utilizing Lipofectamine 3000 (Invitrogen; L3000008). 16 hours later cells were sorted for GFP+ cells and plated. After 32 hours cells were transfected with an equimolar mix of CK2α/β plasmid (Addgene; #27093) and the relevant MCAM tail variant. Following another 16-hour incubation cells were treated with DOX. After 24 hours cells were washed with PBS+EDTA, collected, resuspended in IP Lysis buffer (Pierce; 87787) with the Halt Protease & Phosphatase inhibitor cocktail (Thermo Fisher Scientific, #78440) and allowed to rotate at 4°C for 1 hour. 10ug of rabbit anti-FLAG antibody (Invitrogen; #740001) was added and allowed to incubate overnight at 4°C. The following day, magnetic Protein A/G beads (Pierce; 88802) were added according to the manufacturer’s instructions and samples were rotated at 4°C for 1 hour prior to magnetic separation, washing the A/G beads, and sample preparation for western blot analysis. Westerns were probed for HA-Tag (Invitrogen; #26183) and Myc (Invitrogen; #MA1-980) for CK2 α and β respectively.

### 2.6. Immunohistochemistry

Freshly dissected tissues were fixed overnight at 4°C in 10% neutral buffered formalin (Sigma; HT501128). Fixed tissues were transferred to 70% ethanol for storage at 4 °C. Processing for histology followed standard protocols with paraffin embedding, sectioning at 5 μm thickness, baking for 1 h at 55 °C, and deparaffinizing with CitriSolv (Decon Labs, 89426-268) and rehydration through graded alcohol/water washes. Antigen retrieval was achieved by boiling samples 15min in citrate buffer (10 mM citric acid, 0.05% Tween 20, pH 6.0) prior to overnight primary antibody staining. Slides were mounted in Fluormount-G (Electron Microscopy Sciences). Slides were imaged on a Leica SP8 White light Laser Confocal microscope.

### 2.7. Antibodies

2.7. Antibodies:
| Antibody | Clone | Manufacturer | Catalogue # | Assay | Dilution | RRID |
| --- | --- | --- | --- | --- | --- | --- |
| anti-Akt | 40D4 | Cell Signaling Technology | 2920 | Western Blot | 1:2000 | AB_1147620 |
| anti-phospho-Akt (Ser473) | D9E | Cell Signaling Technology | 4060 | Western Blot | 1:2000 | AB_2315049 |
| anti-β-actin | W16197A | Biolegend | 664802 | Western Blot | 1:2000 | AB_2721349 |
| anti-phospho-Stat3 (Tyr705) | D3A7 | Cell Signaling Technology | 9145 | Western Blot | 1:1000 | AB_2491009 |
| anti-Stat3 | 124H6 | Cell Signaling Technology | 9139 | Western Blot | 1:1000 | AB_331757 |
| anti-Fyn | E3 | Santa Cruz Biotechnology | SC-365913 | Western Blot | 1:500 | AB_10842309 |
| anti-Lyn | C13F9 | Cell Signaling Technology | 2796T | Western Blot | 1:1000 | AB_2138391 |
| anti-phospho-src-family (Tyr418) | EP503Y | Abcam | Ab40660 | Western Blot | 1:1000 | AB_776106 |
| anti-Mcam | 281 | Novus Biologicals | NBP2-90721 | Western Blot | 1:1000 | unavailable |
| anti-MCAM | EPR3208 | Abcam | Ab75769 | Western Blot | 1:1000 | AB_2143375 |
| anti-phospho-Pten (Ser380/Thr382/383) | 44A7 | Cell Signaling Technology | 9549 | Western Blot | 1:1000 | AB_659891 |
| anti-Pten | A2B1 | Santa Cruz Biotechnology | SC-7974 | Western Blot | 1-500 | AB_628187 |
| anti-phospho-Fak (Ty925) | polyclonal | Cell Signaling Technology | 3284 | Western Blot | 1:1000 | AB_10831810 |
| anti-phospho-Fak (Tyr397) | D20B1 | Cell Signaling Technology | 8556 | Western Blot | 1:1000 | AB_10891442 |
| anti-Fak | polyclonal | Cell Signaling Technology | 3285 | Western Blot | 1:1000 | AB_2269034 |
| anti-phospho-Stat5 (Tyr694) | D47E7 | Cell Signaling Technology | 4322 | Western Blot | 1:400 | AB_10544692 |
| anti-Stat5 | ST5-8F7 | Thermo Fisher Scientific | 33-5900 | Western Blot | 1:200 | AB_2533129 |
| anti-Rabbit-IgG-AF680 | polyclonal | Thermo Fisher Scientific | A-21076 | Western Blot | 1:1000 | AB_2535736 |
| anti-Mouse-IgG-AF790 | polyclonal | Thermo Fisher Scientific | A11375 | Western Blot | 1:1000 | AB_2534146 |
| anti-Rat-IgG-H&LDyLight-800 | polyclonal | Thermo Fisher Scientific | SA5-10032 | Western Blot | 1:10000 | AB_2556612 |
| anti-Paxillin | Y113 | Abcam | Ab32084 | Western Blot | 1:5000 | AB_779033 |
| HA-Tag | 2-2.2.14 | Invitrogen | 26183 | Western Blot | 1:1000 | AB_10978021 |
| c-Myc | 9E10 | Invitrogen | MA1-980 | Western Blot | 1:1000 | AB_558470 |
| phospho-CK2 substrate | MultiMab | Cell Signaling Technology | 8738S | Western Blot | 1:1000 | AB_2797653 |
| anti-Sox10 | SD204-04 | Thermo Fisher Scientific | MA5-32398 | Immunocytochemistry | 1:50 | AB_2809676 |
| anti-Krt14 | LL002 | Thermo Fisher Scientific | MA5-11599 | Immunocytochemistry | 1:50 | AB_10982092 |
| anti-Krt8 | TROMA-1 | DSHB U of Iowa | Ab-531826 | Immunocytochemistry | 1:50 | AB_531826 |
| anti-Rabbit-IgG-AF568 | polyclonal | Thermo Fisher Scientific | A-11011 | Immunocytochemistry | 1:200 | AB_143157 |
| anti-Mouse-IgG-AF647 | polyclonal | Thermo Fisher Scientific | A28181 | Immunocytochemistry | 1:200 | AB_2536165 |
| anti-Rat-IgG-AF647 | polyclonal | Thermo Fisher Scientific | A-21247 | Immunocytochemistry | 1:200 | AB_141778 |
| anti-Mcam | ME9F1 | Biolegend | 313605 | FACS | 1:100 | AB_345299 |
| anti-CD326 | G8.8 | Thermo Fisher Scientific | 25-5791-80 | FACS | 1:160 | AB_1724047 |
| anti-Sca1 | D7 | Thermo Fisher Scientific | 17-5981-81 | FACS | 1:300 | AB_469486 |
| anti-Ncam1 | EPR21827 | Abcam | Ab220360 | FACS | 1:500 | AB_2927664 |
| rabbit anti-FLAG | polyclonal | Invitrogen | 740001 | Immunoprecipitation | 1:50 | AB_2610628 |

### 2.8. scRNA Sequencing

All protocols used to generate scRNA-seq data on 10x Genomics Chromium Controller platform including library prep, instrument and sequencing setting can be found on: https://www.10xgenomics.com/support/single-cell-gene-expression/documentation. The Chromium Next GEM Single Cell 3’ Kit v3.1 (PN-1000268) was used to barcode individual cells with 16nt 10x Barcode, to tag cell specific transcript molecules with 12 nt Unique Molecular Identifier (UMI) and to capture poly (A) mRNA with 30 nt ploy(dT) sequence according to the manufactures. The following protocol based on 10x Genomics user guide (CG000315) was performed by High-Throughput Genomics Shared Resource at Huntsman Cancer Institute, University of Utah. Briefly, Py230 single cell suspension was isolated by trypsinization and resuspended in phosphate buffered saline with 0.04% bovine serum albumin. The cell suspension was filtered through 40-micron cell strainer. Viability and cell count were assessed on Countess 2 (Invitrogen, Carlsbad, CA). Equilibrium to targeted cell recovery of 6,000 cells along with 10x Gel Beads and reverse transcription reagents were loaded to Chromium Single Cell Chip G (PN-1000120) to form Gel-Bead-In Emulsions (GEMs), the nano-droplets. Within individual GEMs, barcoded cDNA generated from captured mRNA was synthesized by reverse transcription at the setting of 53°C for 45 min followed by 85°C for 5 min. Subsequent fragmentation, end repair and A-tailing, adaptor ligation and sample indexing with dual index (PN-1000215) were performed in bulk according to the user guide. The resulting barcoded libraries were qualified using Agilent D1000 ScreenTape on Agilent Technology 2200 TapeStation system and quantified by quantification PCR using KAPA Biosystems Library Quantification Kit for Illumine Platforms (KK4842). Multiple libraries were then normalized, pooled and sequenced on NovaSeq 6000 with 150×150 paired end mode.

### 2.9. Bulk RNA-sequencing

RNA was collected from snap frozen biological replicates of 10e6 Py230 shControl/shMcam cell-derived tumors that were collected 12-weeks after inoculation. RNA was isolated via QIAzol-chloroform extraction followed by column-based purification and On-Column DNase Digestion (Qiagen #79254). The aqueous phase was brought to a final concentration of 50% ethanol, and RNA was purified using the RNeasy Lipid Tissue Mini Kit according to the manufacturer’s instructions (Qiagen #74804). Library preparation was performed using the Illumina TruSeq Stranded mRNA Library Prep with UDI (Illumina; poly(A) selection). Sequencing was performed using the NovaSeq 6000 (50 x 50 bp paired-end sequencing; 25 million reads per sample).

### 2.10. Bioinformatic Analysis

All downstream analysis of sequencing data was completed using Loupe Browser (6.0.0) (RRID:SCR_018555), Enrichr (RRID:SCR_001575), Appyters (RRID:SCR_021245), GraphPad Prism (RRID:SCR_002798) and RStudio (4.1.2) (RRID:SCR_000432). For differential gene expression analysis, we utilized the default clustering method of Seurat with the Louvain algorithm to iteratively group cells together ^36^. Reanalysis of primary mouse mammary data from Giraddi et al. used the published diffusion map coordinates and the markers Krt14, Krt8 and Wfdc18, to designate 4 adult groups/cell-types, basal, luminal differentiated, luminal progenitor, and alveolar ^1^. We used the Seurat default differential expressed gene (DEG) test [FindMarkers()] (non-parametric Wilcoxon rank sum test). The top 100 DEGs were then used for downstream comparative analysis. From the Py230 dataset, DEG lists were generated by comparing the shCon luminal to shMcam luminal clusters and similarly for the alveolar clusters identified through marker gene comparisons. We used GSEA software (RRID:SCR_005724), and Molecular Signature Database (MSigDB; RRID:SCR_016863) to compare gene sets with 100,000 permutations in GSEA Pre-ranked. GSEA data was plotted with GraphPad Prism (RRID:SCR_002798). Kaplan Meier plots were generated using TCGA data visualized utilizing KMplot and the auto-select cutoff values for Basal (3.45), Luminal A (3.35), Luminal B (3.29) and Her2 (2.97). ^37^. Heatmaps and MCAM CNV analysis utilized the UCSC Xena Browser ^38^ (RRID:SCR_018938).

### 2.11 Tumor studies

10-week-old nude female mice were from Charles River Laboratories (Wilmington, MA). 1,000 or 1,000,000 MMTV-PyMT Py230 cells were resuspended in complete Matrigel (Corning) and orthotopically injected into the #4 mammary glands of mice. At endpoint, mice were euthanized according to AVMA guidelines and tissues were harvested for processing. The maximum mammary tumor size permitted is 2cm diameter, which was not exceeded in these studies.

### 2.12. Statistics

Replicates and experimental repetition are indicated in figures and figure legends. For statistical analysis, One-Way ANOVA with Tukey Kramer Multiple Comparisons, Repeated Measures One-Way ANOVA (Geisser-Greenhouse correction) w/ Tukey multiple comparison tests, and Two-Way ANOVA with Sidak’s multiple comparisons were used. *\* p<0.05, ** p<0.01, *** p<0.001, **** p<0.0001.* Gene mapping and differential gene expression analysis utilized established default methods embedded in the Cell Ranger, CLoupe browser and Seurat computational packages.

## 3. RESULTS

### 3.1. Mcam maintains epithelial phenotypes in luminal mammary cancer cells

Although experimental OE of MCAM has been reported to drive EMT and basal phenotypes in breast cancer cells ^21,24^, we found that diverse murine/human breast cancer cell lines and patient derived organoid models expressed MCAM including luminal and LP-like models (**Fig. 1A-C**).^9,39^ To test the requirements for Mcam in Luminal breast cancer cell state control and tumorigenicity, we focused our initial attention on the murine Py230 cell line, given its experimental tractability, LP-like phenotype, robust Mcam expression, and reported subtype plasticity.^9,39^ We knocked Mcam down by transient transfection with a panel of Mcam-directed siRNAs (**Figs. S1A-C**) or by stable transduction with lentiviral vectors (LV) bearing an Mcam-directed shRNA construct (**Figs. 1D-H, Fig. S1D-G**). Although transfection efficiency and KD were modest in transient transfections, a subset of cells in siMcam-transfected cultures exhibited qualitative differences in adhesion and cellular morphology with an elongated spindeloid-like phenotype (**Figs. S1A-C**). Although mesenchymal phenotypes were unexpected in KD cells, we obtained corroborating results using sorted stable LV-transduced Mcam KD Py230 cells compared to cells bearing scrambled control vectors (**Figs. 1E-H**). Three independently derived stable Mcam KD clones showed the same cellular and molecular phenotypes observed in transient KD cells including elongated cellular morphology and augmented Paxillin levels (**Figs. 1D-H, S1D-E**). Py230 shMcam cells also exhibited alterations in focal adhesion signaling including elevated activation of focal adhesion kinase and alterations in Src family kinases including Fyn and Lyn (**Fig. 1H**).^28^ These changes were correlated with enhanced migration in scratch wound assays but no significant alteration to proliferation or survival relative to scrambled shRNA controls (**Fig. 1G, S1F**). However, MCAM KD did alter growth kinetics of Py230 in 3D organoid culture as well as the 3D growth of the patient derived organoid model, HCI-011 (**Fig. S1G**).^34^ These data indicate that in contrast to studies where Mcam OE drives mesenchymal phenotypes in epithelial cells, MCAM can also sustain the luminal phenotypes in breast cancer cells.

**Fig. 1.**
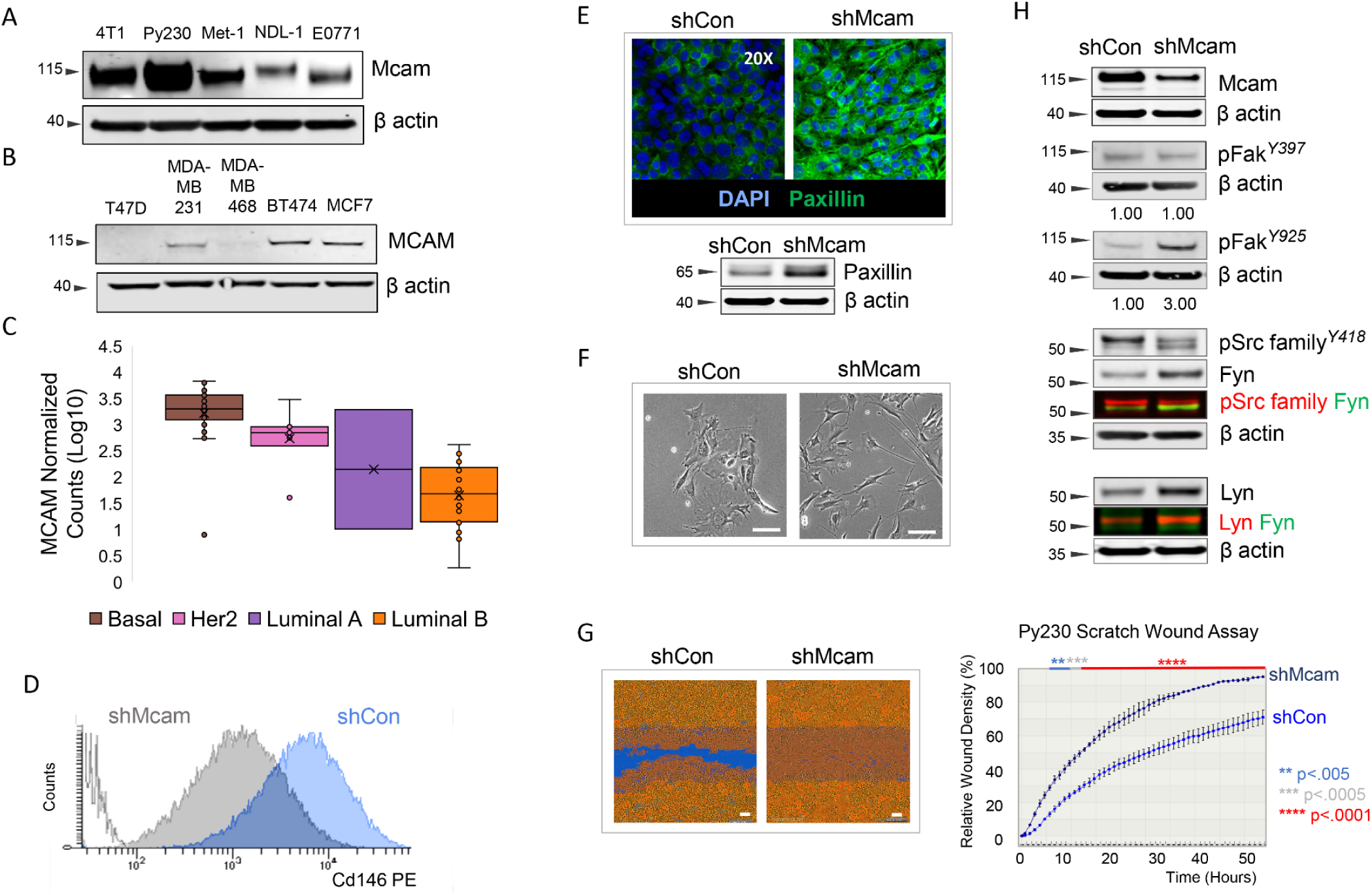
Cellular and molecular signaling changes following Mcam KD in Py230 cells. A) Western blot of mouse breast cancer cell lines. B) Western blot of human breast cancer cell lines. C) Normalized MCAM counts PDM, stratified across breast cancer subtypes by PAM50 analysis. D) Surface Mcam expression (FACS) in control and Mcam KD Py230 cells. E) Representative immuno-fluorescent Paxillin staining and western blot in Mcam KD Py230 cells and scrambled controls. F) Phase contrast image of low density Mcam KD Py230 cells showing a more elongated spindeloid appearance than controls. Scale bar = 100μm. G) IncuCyte scratch wound assay indicates Mcam KD increases migration of Py230 cells. Scale bar = 100μm. Two-way ANOVA with Sidak’s multiple comparisons test between shCon and shMcam. ns = not significant, *p<0.05, **p<0.01, ***p<0.001, ****p<0.0001. H) Western blot analysis of Mcam and altered focal adhesion related signaling in Mcam KD Py230 cells and scrambled controls.

### 3.2. Mcam stabilizes a luminal progenitor cell state in Py230 cells

Changes in cell morphology and morphological heterogeneity were also evident in confluent Mcam KD Py230 cultures relative to controls, suggesting alterations in cellular differentiation states or their relative distributions (**Fig. 2A**). Examination of the lineage related keratins, Krt8 and Krt14, revealed a marked reduction in the frequency of uncommitted Krt8/14 co-expressing cells upon Mcam KD (**Fig. 2B**). To further characterize Mcam-associated cell state changes, we sequenced transcriptomes from several thousand individual control and Mcam KD Py230 cells (**Fig. 2C**). UMAP graphical representation of the data revealed three major transcriptional cell states with differential representation between control and Mcam KD cells (**Fig. 2C, S2A,B**). Surveying the levels of previously described markers of distinct mammary cell states ^2^, we observed that transcript levels for Epcam and Sca1 delineate the major transcriptional states identified in the UMAPs as Epcam+Sca1^High^, Epcam+Sca1^Low^ and Epcam-Sca1^High^. We further determined that the Epcam-cells are Ncam1+ whereas Epcam+ cells are Ncam-(**Fig. 2C**). Flow cytometric analysis (FACS) of surface protein expression mirrored scRNA-seq data and confirmed population skewing upon Mcam KD (**Fig. 2D, S2C,D**). FACS and scRNA-seq of additional independent clones demonstrated consistent loss of the Epcam+Sca1^High^ cell state in Mcam KD cells, although some variance in the final population frequencies was observed (**Figs. S2B-D**). Upon Mcam OE with a silent mutation-bearing shRNA-resistant vector in our Py230 Mcam KD cells, we restored predominance of the Epcam+Sca1^High^ profile while Epcam+Sca1^low^ and Epcam-Ncam+ cell populations were restored to near baseline proportions (**Figs. S2E-H**). We also determined that all three populations were able to interconvert and re-establish parental distributions within 3 weeks of cell sorting by FACS, although kinetics for redistribution from the Epcam-Ncam+ of each genotype were slower than other populations (**Fig. 2E**). Critically, while sorted Epcam+Sca1^Low^ cells of both genotypes rapidly (within one week) generated Epcam+Sca1^High^ cells, the Epcam+Sca1^High^ cell type was depleted in Mcam KD cells within two weeks (**Fig. 2E**, arrows). Thus, Mcam KD alters cell state distributions in Py230 by destabilizing the Epcam+Sca1^High^ phenotype.

**Fig. 2.**
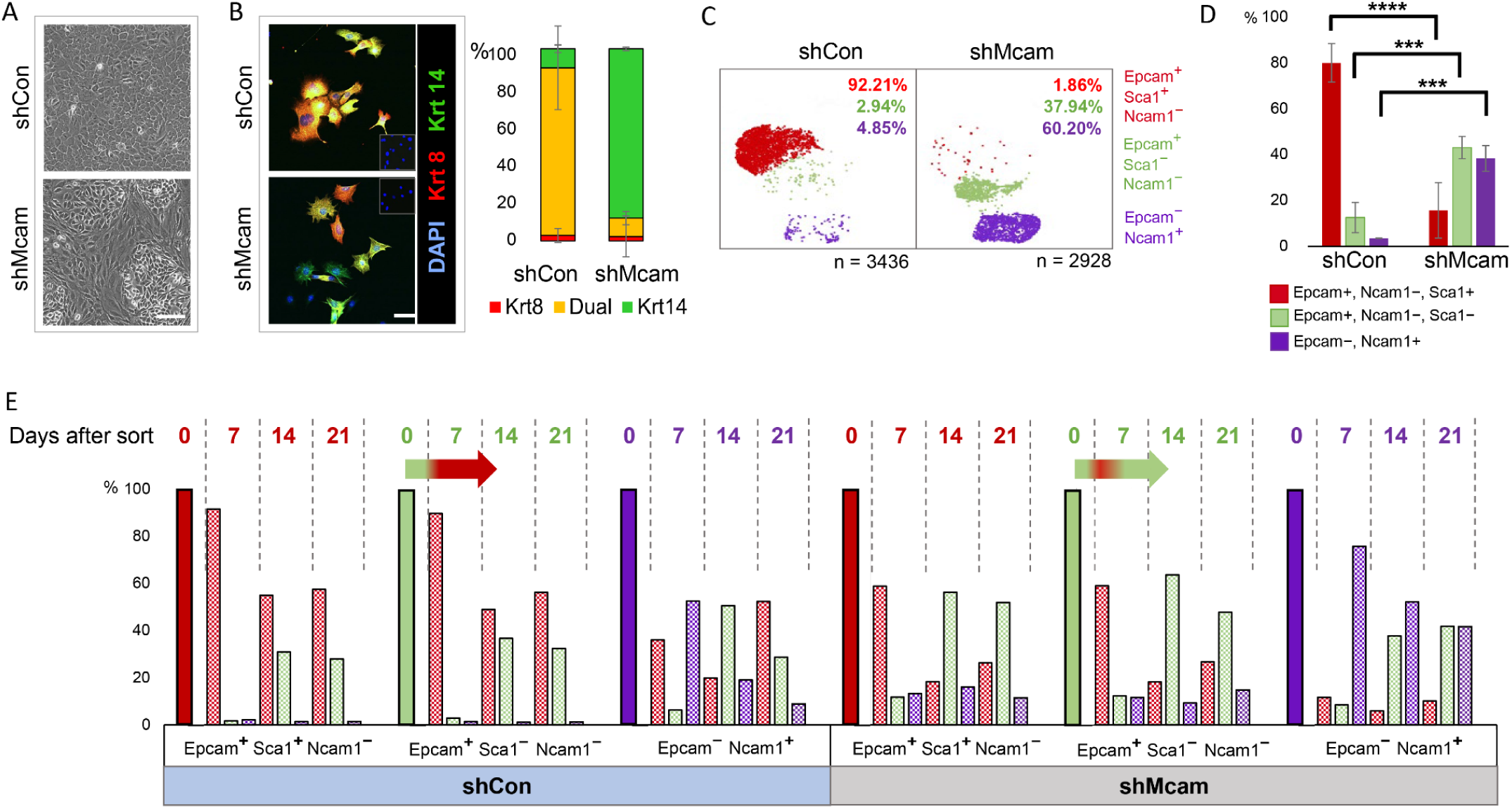
Cell-state change following Mcam KD in Py230 cells. A) Phase contrast image of confluent Mcam KD and control Py230 cultures. Scale bar = 100μm. B) Altered expression of lineage associated Krt8 (red) and Krt14 (green) in Py230 Mcam KD cultures. Scale bar = 100μm. C) scRNA-seq profiles from control and Mcam KD cells reveal three major subpopulations that are identifiable by expression of previously described mammary lineage markers ^2^. D) FACS analysis in triplicates confirms proportional skewing among cells expressing these three marker proteins that mirrored subpopulations identified by scRNA-seq (n=3 independent clones per genotype). Error bars = SD. Two-way ANOVA with Tukey multiple comparison test: ns = not significant, *p<0.05, **p<0.01, ***p<0.001, ****p<0.0001. E) Interconversion of Py230 subpopulations upon FACS separation and culture for 0, 7, 14 or 21 days. Horizontal arrows show immediate conversion of Epcam+Sca-Ncam-cells to Sca+ within seven days followed by reversion in to Sca-predominance in Mcam KD cells by day 14.

### 3.3. Mcam loss triggers a Stat3 driven alveolar/basal lineage switch

Epcam and Sca1 have previously been proposed to distinguish alveolar progenitors (AP; Epcam^high^Sca1^low^) from hormone sensing luminal progenitors (HSP/LP; Epcam^high^Sca1^high^).^2^ We took a comparative transcriptomics approach to determine whether Py230 transcriptomes mimic these progenitor states more broadly. We determined that Py230 cells shared expression similarity with previously published expression profiles for HSP/LP transcriptional states, while the majority of Mcam KD Py230 cells resembled AP and basal cell states of the normal gland (**Figs. 3A-C, S3A-C**).^1,40^ Additionally, although we noted upregulation of genes previously ascribed to neural crest fates in Mcam KD Py230 cells *in vivo*, we determined that several of these genes are also expressed in normal mammary alveolar and basal cell types and do not necessarily indicate loss of mammary epithelial specification (**Fig. 3C, S3A**).^1,40,41^

**Fig. 3.**
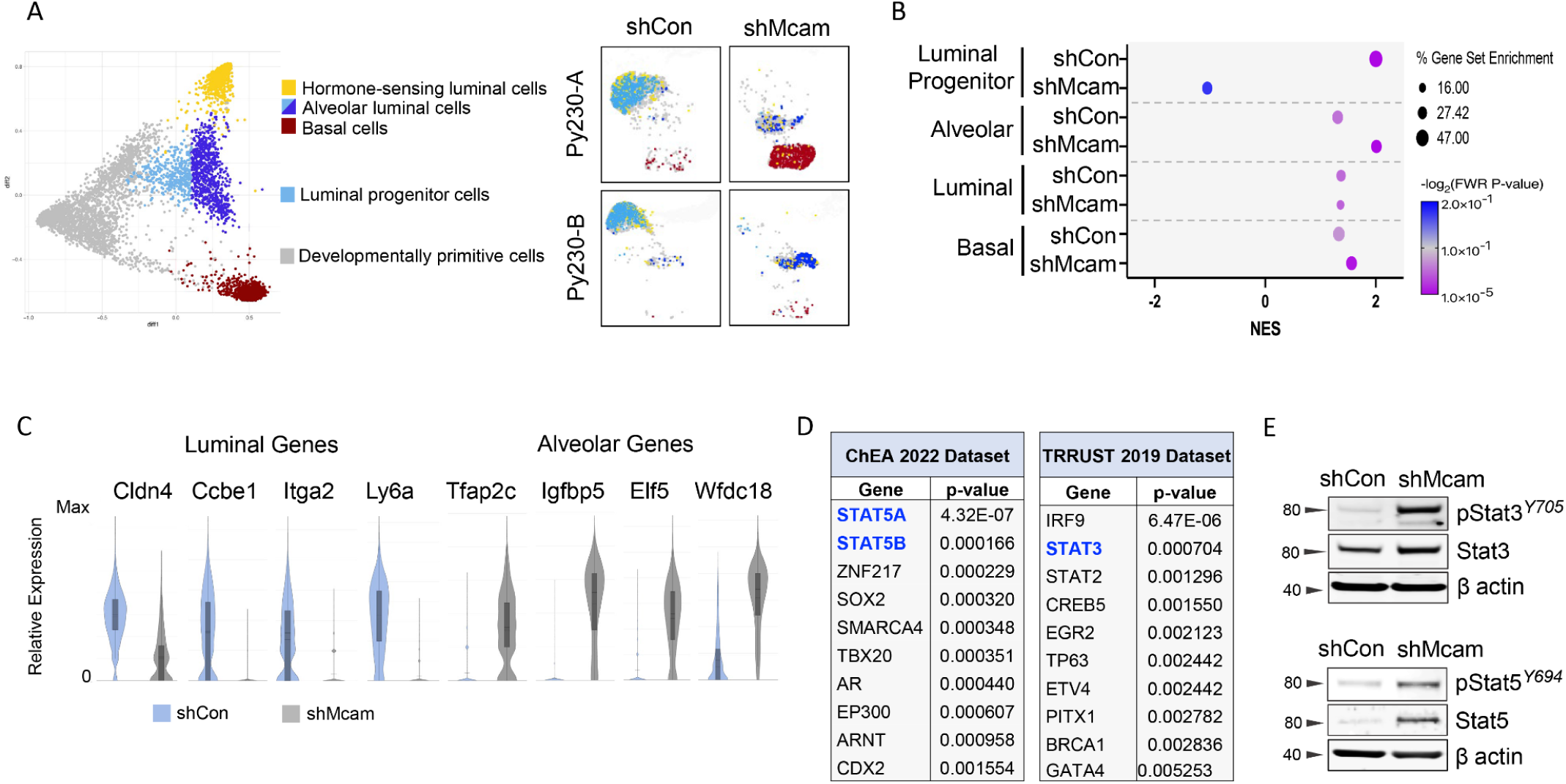
A luminal progenitor to alveolar/basal lineage switch accompanies MCAM loss in Py230 cells. A) The top and bottom five differentially expressed genes between the adult mammary cell types identified in the diffusion map from Giraddi et al. (left) ^1^,identify different cell states in Py230 control and Mcam KD cells (right). B) Gene set enrichment analysis (GSEA) showing enrichment of shCon and shMcam for luminal progenitor and alveolar cell states, respectively compared to normal mouse mammary cell populations from Giraddi et al. ^1^. C) scRNA-seq analysis for relative expression of luminal and alveolar cell state markers in Py230 control (blue) and Mcam KD (grey) cells. D) Top 10 genes associated with shMcam Py230 cells from Enrichr analysis using ChEA and TRRUST datasets. E) OE and phosphorylated Stat3 and Stat5 in Py230-A cells.

To identify underlying regulatory programs contributing to MCAM mediated cell state control, we examined enriched transcription factor target categories among genes that were differentially expressed between control and Mcam KD Py230 cells using EnrichR.^42^ In addition to Elf5, a classic alveolar lineage specifier (**Fig. 3C, S3B**)^43^, genes upregulated in Mcam KD cells were significantly enriched for target genes of the STAT family of transcription factors, including alveolar markers Tfap2c, Igfbp5, and Wfdc18 (**Figs. 3C,D**).^17,18,44–46^ We therefore examined Stat3 and Stat5a as potential mediators of Py230 cell state change consequent to Mcam KD. Activated Stat3 and Stat5A were upregulated in Mcam KD Py230, providing a potential mechanistic basis for the alveolar cell state switch adopted by Mcam KD cells and for the OE of Stat target genes described above (**Fig. 3E, S3D-F**). While increased Stat5a could be explained by its overexpression, total Stat3 protein levels were only modestly upregulated though it was strongly overactivated in Mcam KD cells relative to MCAM expressing controls (**Fig. 3E, S3D,E**).

### 3.4. Mcam governs Ck2 substrate utilization attenuating Stat3 activity in Py230 cells

To determine whether hyper-activation of Stat3 was necessary for the HSP/LP ◊ AP switch, we treated Py230 cells with Stattic, a selective STAT3 inhibitor. Additionally, we looked at the effects of inhibitors of upstream regulators of STAT signaling including Ag490, an inhibitor of JAK2 and EGFR, and the CK2 inhibitors CX-4935 and GO289, (**Figs. 4A-E, 4G, S4A-D**).^44,47,48^ Levels of phosphorylated Stat3 (pStat3) in Py230 Mcam KD cells were decreased to near baseline levels with Stattic or by inhibition of either upstream kinase (**Fig. 4A**). Cell state skewing associated with Mcam KD was dramatically reversed by CX-4945, and to a lesser degree by Stattic, whereas effects of Ag490 were not significant with respect to restoration of a predominant Epcam+Sca1^High^ LP-like cell state (**Figs. 4B-E, S4A,B,E-H**). Cytotoxicity was negligible for all three reagents. The role of Ck2 in Mcam KD phenotypes was intriguing as CK2 has not previously been implicated as a functional downstream target of MCAM, although CK2 has previously been implicated in breast tumorigenesis and cell state control of mammary cells.^49,50^ In previous studies, reduction of CK2 levels in the colorectal cancer cell line, LoVo, led to reduced E-cadherin and other signs of EMT.^51^ However, in our study, Ck2 inhibition did not significantly alter EMT features and Ck2 levels were unaffected by Mcam KD (**Figs. S4D,H**). Rather, we noted Mcam KD was associated with augmented Ck2 activity toward Stat3 and phosphorylation of the PI3K negative regulator Pten at Ck2 specific sites (Ser380/Thr382/Thr383) that promote its degradation and consequent Akt hyperactivation (**Fig. 4F, S4I**).^51,52^ Ck2 inhibition reversed these effects and restored expression of luminal markers while depleting alveolar markers (**Fig. 4G, S4D**). These data indicate that Mcam controls cell state in Py230 cells in a manner dependent on Ck2 and Stat3.

**Fig. 4.**
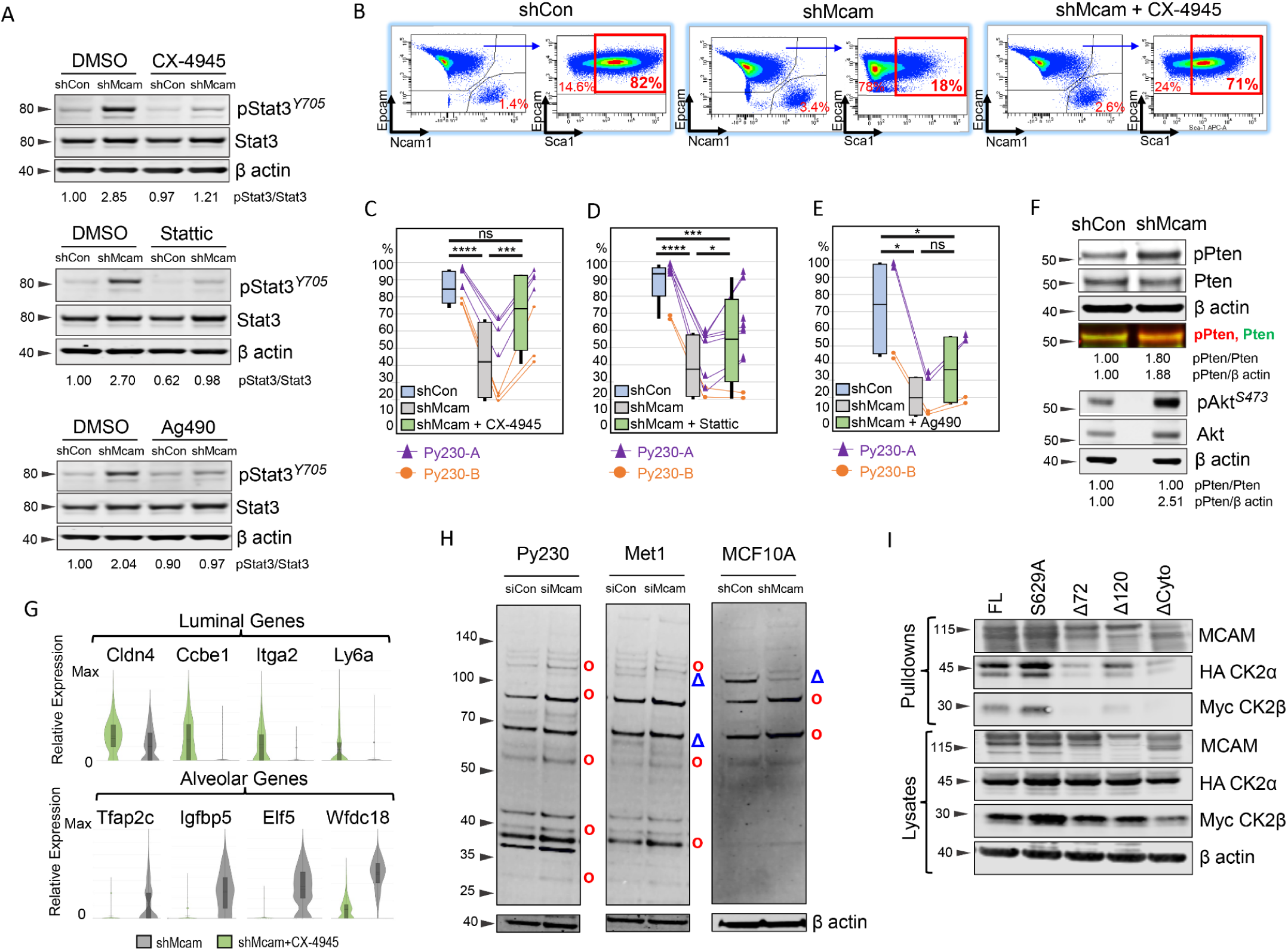
Overactive CK2 in Py230 shMcam cells maintains the alveolar progenitor phenotype. A) Over-expressed and phosphorylated Stat3 in Py230-A cells and its inhibition by the kinase inhibitors, CX-4945 10μM, Ag490 50μM, Stattic 1μM for 30 minutes in WB analysis. B) Representative FACS analysis of Py230-B cells treated with CX-4945 10μM for 7 Days and stained for Epcam, Sca1 and Ncam1. C-E) Compiled FACS analysis showing the rescue of Epcam+, Sca1+ subpopulation frequencies in Py230-A and Py230-B cells treated with CX-4945 10μM for 7 Days (C), with Stattic 1μM for 7 days (D), and with Ag490 30μM for 7 days (E). RM one-way ANOVA (Geisser-Greenhouse correction) with Tukey multiple comparison test: *p<0.05, **p<0.01, ***p<0.001, ****p<0.0001. F) Western blot analysis in Py230-A cells showing higher basal levels of pPten (Ser380/Thr382/383) and pAKT in Mcam KD Py230. G) Gene expression in Py230-A cells treated with CX-4945 10μM for 7 Days (scRNA-seq). H) Western blot analysis in select breast cancer cell lines showing differential protein expression across phosphorylated CK2 substrates in siCon and siMcam transfected cell lines in culture for 48 hours post transfection or with shCon and shMCAM transduction. o=increased in MCAM KD, Δ=decreased in MCAM KD. I) Co-immunoprecipitation western blot analysis of a transiently co-transfected Flag-tagged Mcam variants and HA-Ck2a/Myc-Ck2b expression constructs in HEK293A cells. FL=full length Mcam, S629A=point mutant in the CK2 phosphorylation site of Mcam, Δ72=deletion of the distal 72 amino acids of the Mcam tail, Δ120=deletion of the membrane proximal 120 amino acids of the Mcam tail, ΔCyto=complete deletion of the cytoplasmic tail of Mcam.

Using an antibody that detects phosphorylated Ck2 (pCK2) sites across diverse proteins, we observed altered Ck2 phosphorylation target patterns as a function of MCAM status in a variety of lines, including Py230, another PyMT derived line, Met1, and the human non-transformed epithelial cells MCF10A (**Fig. 4H, S4J,K**). CX-4945 treatment confirmed Ck2 specificity of the majority of detected proteins (**Fig. S4L**). Further, Flag ‘pull downs’ of Flag-tagged MCAM rescue constructs co-precipitated tagged CK2α/β proteins when co-expressed in HEK293A cells, indicating Mcam regulation of Ck2 may be direct and dependent on the terminal 72 amino acids of MCAMs’ cytoplasmic (**Fig. 4I**). Mutation of a putative Ck2 phosphorylation site target residue (S629A) did not compromise binding (**Fig. 4I**).^31^

### 3.5. MCAM loss hallmarks aggressive luminal tumors

Considering the MCAM expression we observed in luminal tumor cell lines and its activity in HSP/LP/AP cell state transitions in Mcam KD Py230 cells, we re-examined MCAM’s relationship to tumor aggressiveness and subtype using The Cancer Genome Atlas (TCGA) database. While we confirmed that elevated MCAM levels correlated with poorer prognosis in Basal-like and Her2-enriched subtypes, the inverse was true in a subset of luminal cell types when examining patients treated with chemotherapy (**Fig. 5A**). Chemotherapy treated luminal cancers tend to reflect more aggressive tumors than those treated with endocrine therapy as an adjuvant monotherapy. Examining MCAM expression across the various breast cancer subtypes, we found that MCAM was generally reduced across all breast cancer subtypes relative to normal samples, yet more highly expressed in Basal-like, and Her2-enriched tumors than in Luminal A/B tumors as previously reported (**Fig. 5B, S5A**).^22^ It should be noted that to some extent MCAM expression levels in TCGA may also reflect different levels of stromal involvement since stromal cells can express high levels of MCAM.

**Fig. 5.**
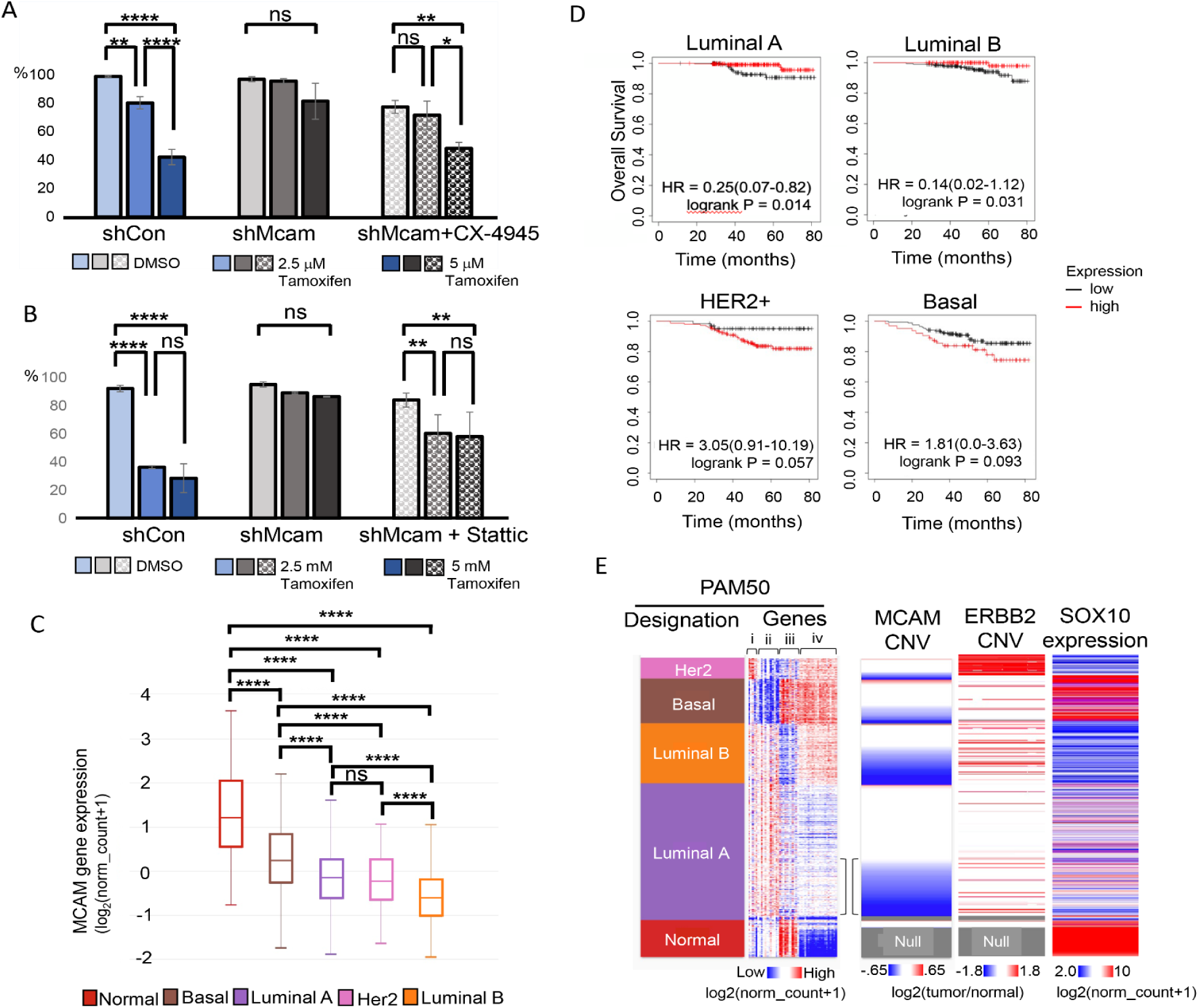
MCAM loss marks aggressive luminal tumors. A-B) IncuCyte proliferation assays of Py230 control and Mcam KD cells with tamoxifen treatment shows that pretreatment with CX-4945 10μM for 7 days (A), and pretreatment with Stattic 1μM for 7 day (B) rescue the tamoxifen sensitivity in Mcam KD cells. One-Way ANOVA with Tukey Kramer Multiple Comparisons. *\* p<0.05, ** p<0.01, *** p<0.001, **** p<0.0001*. C) MCAM expression across breast cancer subtypes in TCGA data. One-Way ANOVA with Tukey Kramer Multiple Comparisons. *\* p<0.05, ** p<0.01, *** p<0.001, **** p<0.0001;* n=1247. D) Kaplan-Meier analysis of TCGA data breast cancer data, stratifying patients receiving chemotherapy between high and low MCAM expression. E) TCGA breast cancer data organized by PAM50 designation and MCAM copy number variation (CNV) with expression of PAM50 gene set, ERBB2 copy number variation, and SOX10 expression. The PAM 50 gene set can be broken down roughly into groups of genes related to ERBB2 expression (i), luminal genes including hormone receptor expression (ii), basal genes (iii), and proliferation genes (iv). Brackets highlight ‘transitional’ Luminal A with increased proliferation and MCAM CNV loss.

The Luminal B intrinsic subtype is distinguishable from Luminal A tumors by its poorer prognosis, more frequent endocrine resistance, and greater proliferation.^3,6^ However, underlying mechanisms that give rise to these distinctions remain unclear. We noted that reduced MCAM gene copy number was surprisingly frequent in breast cancer, especially in luminal tumors (**Fig. 5C, S5A**). In the Luminal A subtype MCAM loss was associated with a trend toward increased expression of proliferation associated PAM50 genes and therefore a trend toward a more Luminal B like phenotype (**Fig. 5C, S5B**).^7,38,53^ This likely reflects concomitant amplification of Cyclin D1 at 11q, in many cases.^53^ Interestingly, among patient derived models (PDMs) that formed estrogen independent outgrowths, two out of four instances showed concomitant reduction in MCAM, one of which (HCI-040) showed evidence of the same 11q perturbations observed in TCGA (**Fig. S5C**).^34^ We also noted a correlation between MCAM copy number reduction and ‘triple positive’ (TPBC) status in archival TCGA data (**Fig. 5C**). Clinically, despite their amplified ERBB2, such tumors are usually treated as HR+ luminal tumors (**Fig. 1C, S5D**).

Frontline treatments for luminal, HR+ breast cancers include a variety of estrogen blocking therapies such as the selective estrogen receptor modulator, tamoxifen. When grown in media containing estrogens, Py230 cells express estrogen receptor and become estrogen sensitive.^9^ In light of this, our observation that Stat3 inhibition reverts Mcam KD Py230 cells to an HSP/LP-related transcriptional state, and prior reports that inhibition of Stat3 can increase the sensitivity to tamoxifen treatment, we tested the sensitivity of control and Mcam KD Py230 cells to tamoxifen using an *in vitro* proliferation assay (**Figs. 5D,E**). We found control cells are more sensitive to tamoxifen than Mcam KD cells, and that sensitivity can be restored in Mcam KD Py230 cells by treatment with inhibitors of Stat3 or Ck2 (**Figs. 5D,E**). Together, these data further indicate that Mcam maintains a hormone-sensing cell state through modulation of Ck2 and downstream Stat3 signaling.

### 3.6. MCAM promotes tumorigenicity through cell state control

Finally, we examined whether cell state and signaling changes observed in Mcam KD cells *in vitro* alter tumorigenic potential of the cells or whether they alter the neural crest associated, Basal-like, Sox10-positive phenotype previously associated with Py230 cells *in vivo* (**Fig. 5C**).^39^ Surprisingly, given the apparent increase in mesenchymal traits observed when Mcam is KD in these cells in 2D culture, they were less tumorigenic than control cells in the graft setting when low cell numbers (1,000) were injected (**Fig. 6A**). When 1,000,000 cells were orthotopically transplanted into the mammary fat pads, we saw formation of both control and Mcam KD-derived tumors, but the tumors that form were fundamentally different in terms of size, histopathological appearance, and expression of select cell type markers (**Fig. 6B**). We found that Py230 cells exhibit expression of Sox10 *in vivo* though they lack such expression in 2D culture, consistent with a prior report (**Fig. 6B**).^39^ Sox10 expression *in vivo* marked large invasive peripheral regions of Py230 tumors consistent with the previously proposed neural crest like differentiation state. We determined cells in such regions co-express Krt14 suggesting they are likely basal-like or luminal progenitor-like regions (**Figs. 5C, 6B**). However, these large uniform Sox10+Krt14+ positive regions were absent in Mcam KD tumors and the many fewer Sox10+ cells that were found in Mcam KD tumors were usually either Krt8/14 negative or were found in organized Krt8+ epithelial structures resembling the Sox10+,HR-luminal cells previously described in the normal mouse mammary epithelium (**Fig. 6B**).^41^ Gene expression profiling of control and Mcam KD tumors confirmed cell state skewing by Mcam KD that was consistent with a block to the luminal progenitor related basal-like phenotype (**Fig. 6C**).

**Fig. 6.**
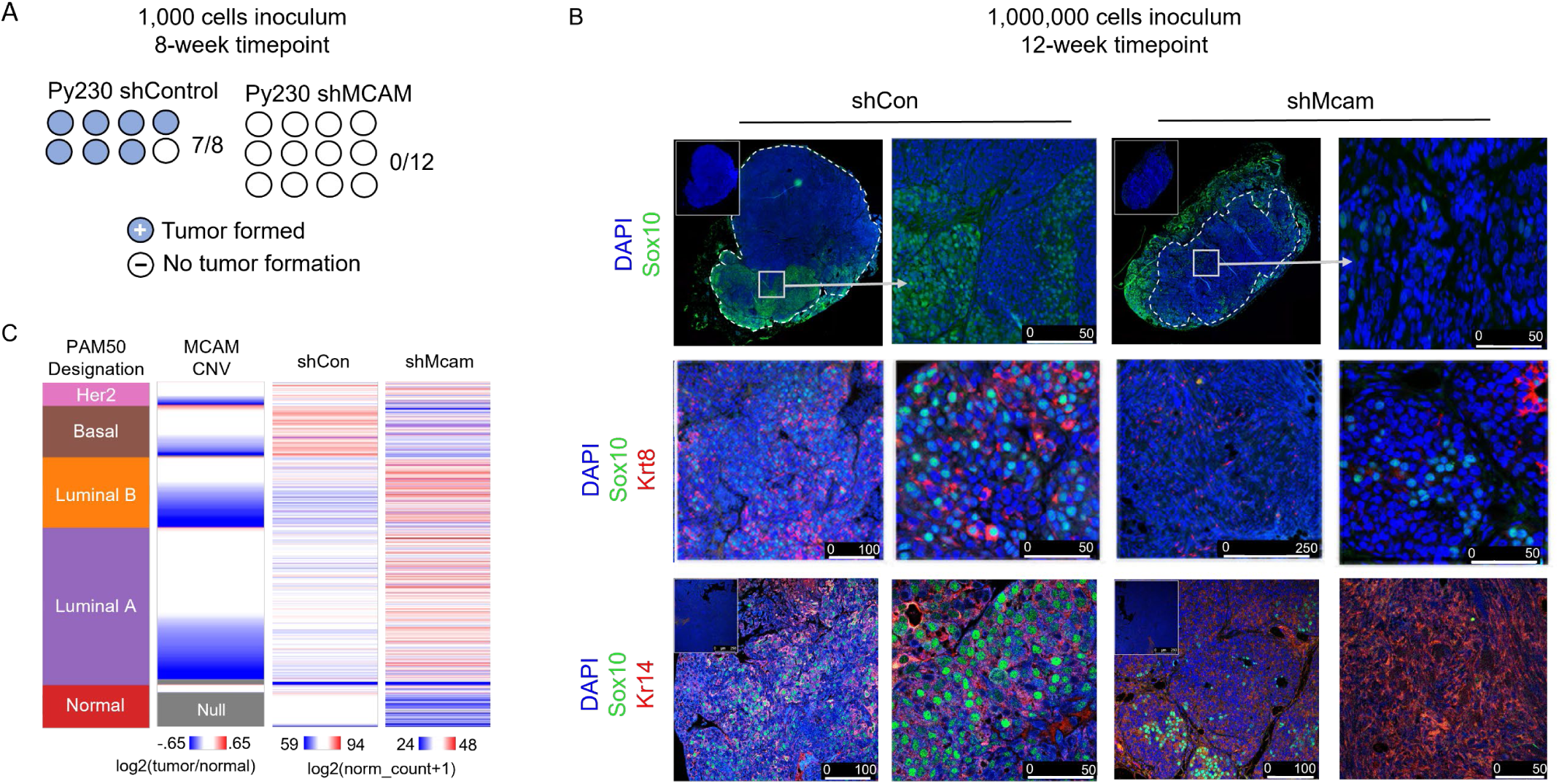
Mcam promotes tumorigenic phenotypes and tumor cell plasticity. A) Reduced tumor seeding capacity of Mcam KD Py230 cells (1,000 cell orthotopic injection) after 8 weeks of growth. B) Altered cell type distribution in tumors *in vivo* generated from orthotopic injection of 10e6 Py230 control or Mcam KD cells. IHC analysis shows differential expression and distribution of cell type markers between control and Mcam KD tumors. Sox10+ stromal tumor margins are demarcated from tumor parenchyma (dashed lines). Regions of interest in low magnification images marked by white squares. C) TCGA breast cancer data organized by PAM50 designation, MCAM copy number loss, and relative expression of genes with ten greatest rank changes up or down between control and shMCAM Py230 tumor grafts (bulk RNASeq). Two-way ANOVA *p<.0001*.

## 4. DISCUSSION

Lineage plasticity is critical for proper mammary gland development and maintenance, however, in the context of breast cancer, lineage plasticity could drive increased tumor heterogeneity and treatment failure ^10,11^. Although OE of Mcam has previously been shown to promote EMT in mammary epithelial cells, here we report significant Mcam expression among luminal epithelial breast cancer cell lines and the surprising finding that Mcam is required to maintain the epithelial LP-like ground state of the Py230 model. This finding is consistent with a recent study identifying MCAM as a marker of KRT14/KRT19 double positive LP cells of the human breast ^54^. Mcam KD in Py230 cells promoted differentiation toward AP/Basal phenotypes with associated alterations in adhesion and motility, as well as insensitivity to tamoxifen and an apparent block to HSP-like and Sox10-positive neural crest-like/basal-like breast cancer cell states.

Mcam effects and altered Ck2 substrate utilization while present across the different cell lines we examined, appear to be cell type and breast cancer subtype specific as 1) expression in the basal-like 4T1 cell line promotes, rather than inhibits, Stat3 activation, 2) several cell lines showed marginal to undetectable Stat3 alterations under Mcam KD, and 3) MCAM expression among HR+ breast cancer PDMs was enriched among those that (like Py230) co-express ERBB2 and classify as Luminal B (**Fig. S5D** and ^55^). We propose that the cell type specificity of MCAM effects may reflect coupling or decoupling of CK2 (or other upstream kinases) to different client proteins depending on a cell’s ground state. This may be critical to understanding CK2 activity and regulation as CK2 is a constitutively active high-level regulatory kinase with hundreds of substrates whose mechanism of regulation and roles in mammary lineage control are not well understood, though scaffolding (for instance by MCAM) represents a plausible mechanism.

Despite the increased mesenchymal features and therapy resistance observed for Mcam KD Py230 cells *in vitro*, Mcam KD reduced tumorigenic seeding and growth of grafted cells in mice, a result that is at least superficially consistent with prior studies correlating higher MCAM expression to tumor aggressiveness.^19,22,24^ However, this too may be subtype dependent as re-analysis of archival TCGA breast cancer data showed MCAM expression to be a favorable marker in Luminal A and B tumors and showed MCAM loss to typify the more aggressive and more frequently tamoxifen resistant Luminal B subtype as well as the more proliferative (by gene signature) samples among Luminal A tumors.^38^ By contrast, tumors robustly expressing MCAM are typically Basal-like. It is noteworthy that recent studies suggest that the Basal-like subtype arises from LPs and ‘involution mimicry’, thus invoking a critical role for LP and AVP fate switches ^43,56^. Based on our data and analysis of TCGA, we speculate that MCAM may be essential for the emergence of Basal-like tumors arising from the LP.

It should be noted that comparative transcriptomics did not indicate a strong association between Luminal B tumors in TCGA and Mcam KD Py230. The recurrent genetic aberration at 11q that typifies most of the TCGA breast cancers with Mcam copy number reduction usually involves concomitant loss of several additional 11q genes that may contribute to cell phenotype (e.g., PGR, ATM, etc.) and amplification of Cyclin D1 (**Figs. S5C,D**).^4,6,53^ Thus, it is reasonable to assume that several additional genetic alterations may cooperate with MCAM loss in the context of Luminal B tumors in humans. Nevertheless, altered tamoxifen responses have been associated with 11q deletions in human breast cancer although the specific gene(s) responsible for this effect remain uncertain.^53^

The findings above may be key to designing combination therapies that effectively manage shifting therapeutic vulnerabilities deriving from the plastic cell state change potential of mammary luminal progenitors. For instance, newer CK2 or STAT3 inhibitors may be of interest in combination with hormone receptor targeted therapies for some cancers.^49,57^ STATs and associated factors are not only strongly implicated in alveolar cell state control in the normal gland but also have well documented roles in breast and other cancers.^18,55,57^ Indeed, a recent study demonstrated STAT3 inhibition can increase the tamoxifen sensitivity of tamoxifen-resistant breast cancer cells.^57^ Alternatively, if MCAM expression permits access to invasive neural crest like states *in vivo,* it may be more effective to inhibit MCAM and seek out new vulnerabilities of the resultant hormone-insensitive cell state. Altogether, our studies indicate that Mcam plays a critical role in breast cancer cell state determination via Ck2 and Stat3 control, with implications for understanding developmental plasticity and MCAM’s role in breast cancer intra- and inter-tumoral heterogeneity.

## 5. DECLARATIONS

## Ethics approval and consent to participate

All animal care and procedures were approved and monitored by an Institutional Animal Care and Use Committee.

## Consent for publication

Not applicable.

## Data Availability

Datasets generated during and/or analyzed during the current study are either publicly available (TCGA; https://www.cancer.gov/tcga), are deposited at Gene Expression Omnibus (GEO) under GSE233093 or are within the article and its supplementary data files. All codes and R-packages used in the study are publicly available and have been disclosed in Methods or are available from the corresponding authors on reasonable request.

## Authors’ Contributions

Concept: BTS. Experimental Design: OB, BLG, BTS. Execution: OB, BLG, DWF, BH, BTS. Analysis: BLG, EMM, DA-T. Interpretation: All authors. Writing: OB, BLG, BTS.

## Acknowledgements

We thank C. Trejo for technical assistance and M. Kruithof-de Julio, T. Oliver, and D. Salomon for valuable discussions. We also acknowledge generous support from the Huntsman Cancer Foundation, and from the Halt Cancer at X foundation. We thank the University of Utah/HCI Cancer Center Support Grant (P30 CA42014), and staff in the Flow Cytometry, HSC Cell Imaging, Preclinical Research Resource, Biostatistics, High-Throughput Genomics and Bioinformatics shared core resources. Research reported in this publication utilized the Preclinical Research Resource, Flow Cytometry Core, Biorepository and Molecular Pathology, and High-Throughput Genomics and Bioinformatics shared resources at Huntsman Cancer Institute at the University of Utah and was supported by the National Cancer Institute of the National Institutes of Health under Award Number P30CA042014. The content is solely the responsibility of the authors and does not necessarily represent the official views of the NIH.

## LIST OF ABBREVIATIONS

TCGA: The Cancer Genome Atlas
scRNA-seq: Single cell RNA-seq
RNA-seq: RNA sequencing
KD: Knock down
OE: Over expression
shRNA: Short hairpin RNA
siRNA: Small Interfering RNA
IHC: Immunohistochemistry
pCK2: Phosphorylated casein kinase 2
GSEA: Gene Set Enrichment Analysis
EMT: Epithelial-to-Mesenchymal Transition
LP: Luminal progenitor
HSP: Hormone-sensing progenitor
AP: Alveolar progenitor
TPBC: Triple Positive Breast Cancer
TNBC: Triple Negative Breast Cancer
CNV: Copy Number Variation
ns: not significant
HR: Hormone Receptor
FDR: False discovery rate
NES: Normalized enrichment score
FACS: Fluorescence-activated cell sorting
ERM: ezrin-radixin-moesin
MMTV-PyMT: Mouse mammary tumor virus polyomavirus middle T
PDM: Patient Derived Model

