## Supplementary Figures for "Mcam stabilizes luminal progenitor breast cancer phenotypes via Ck2 control and Src/Akt/Stat3 attenuation"

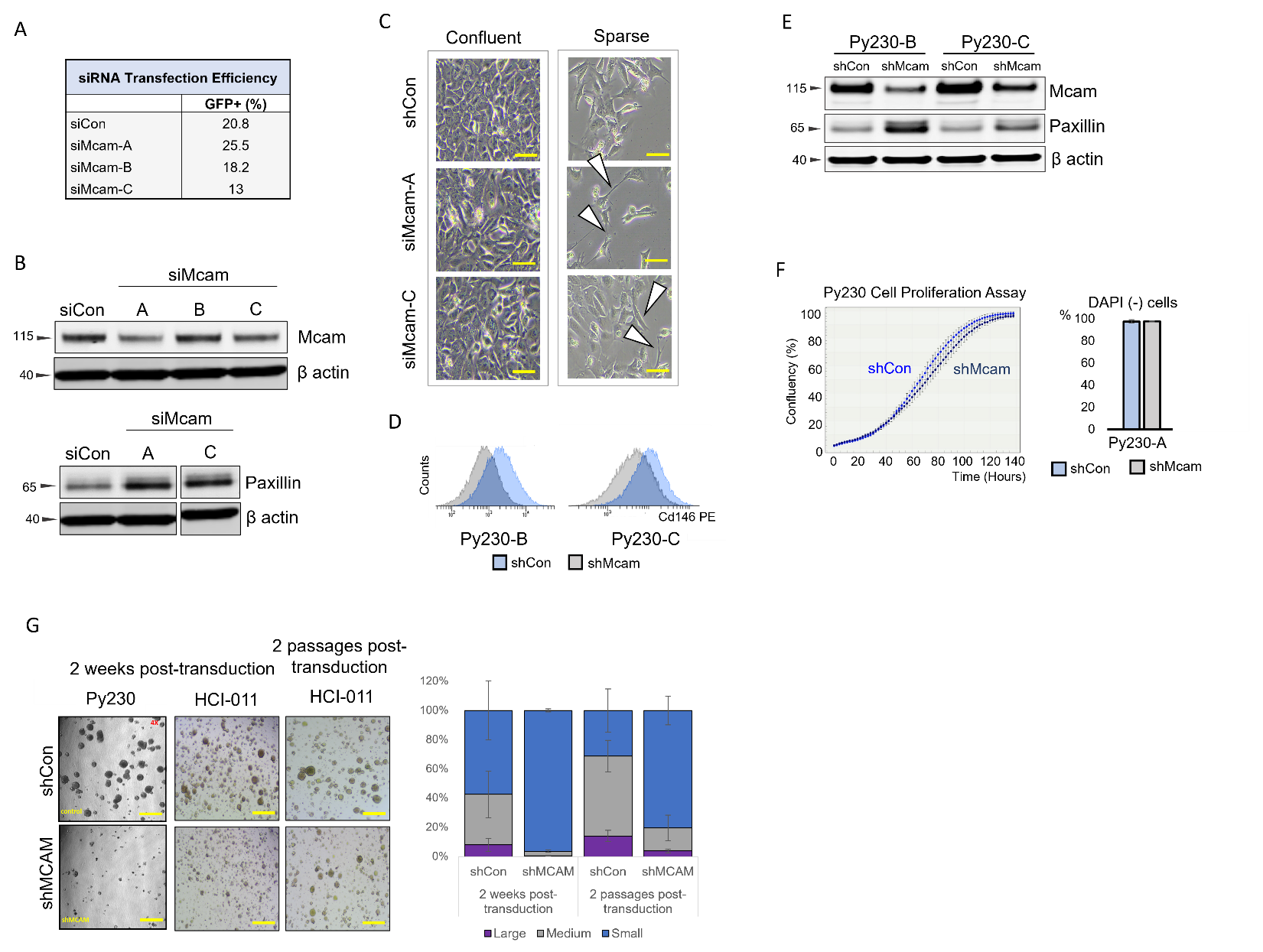

**Supplemental Figure 1. Mcam expression across murine mammary cancer cell lines and consistent cellular and molecular features in Mcam KD Py230 cells.** A) Quantification of flow cytometric analysis of relative KD efficiency by GFP expression across multiple siRNA knockdown clones and scrambled control (siCon) in Py230 cells. Unshaded histograms indicate unstained controls. B) Western blot analysis of independent transient transfection of siMcam in Py230 cells probed for Mcam (clones A-C) and Paxillin (clones A and C). C) Phase contrast image of confluent and sparsely plated siMcam clones A and C and control Py230 cultures. White arrows pointing towards elongated structures of interest. D) Reduced overall Mcam levels in independent clones of Mcam KD Py230 by flow cytometric analysis. E) Mcam KD consistently increases total paxillin levels in independent Mcam KD Py230 clones. F) Survival ratios from DAPI analysis with fluorescent cytometry in different clones of Py230 cells. G) Phase contrast imaging of stably transduced Py230 organoids and the patient derived HCI-011 line at 2 weeks post transduction and HCI-011 at 2 passages post-transduction showing a return to the control phenotype after 2 passages in HCI-011 cells. Scale bar=200μM. Right panel is quantification of HCI-011 organoid size at 2 weeks post-transduction and 2 passages post-transduction.

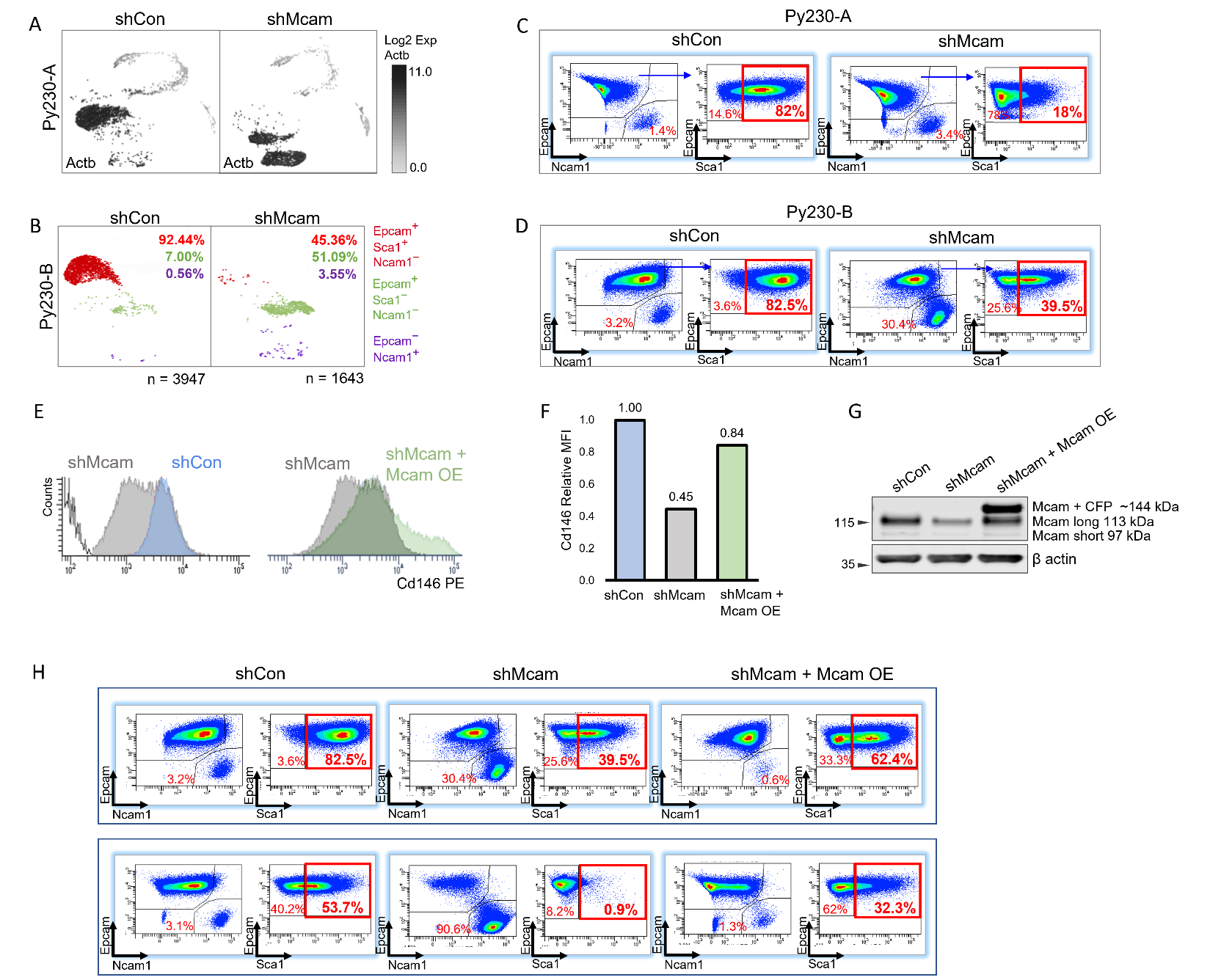
**Supplemental Figure 2. Cell-state change following Mcam KD in Py230 cells and Mcam rescue with an Mcam overexpressing vector in Mcam KD Py230 cells.** A) Unsupervised graphing of scRNA-seq profiles with Actb from control and Mcam KD cells reveals three major subpopulations. B) scRNA-seq of Py230-B clone demonstrates consistent loss of the Epcam+Sca1^High^ cell state when Mcam was knocked down. Percentages refer to the total overall percentage of cells in each population. C) Representative flow cytometric analysis in Py230-A and D) in Py230-B cells showing very similar profile to that of flow cytometry analysis in terms of subpopulations. E) FACS analysis of Py230-A control, Mcam KD and Mcam KD cells transduced with an Mcam overexpression vector. Unshaded histogram indicates unstained control. F) Bar graphs showing the Median Fluorescent Intensities for Mcam in cells from E. G) Western Blot Analysis for rescued Py230-A Mcam KD cells. H) FACS analysis shows that the subpopulation frequencies are partially restored in rescued Mcam KD cells (n=2).

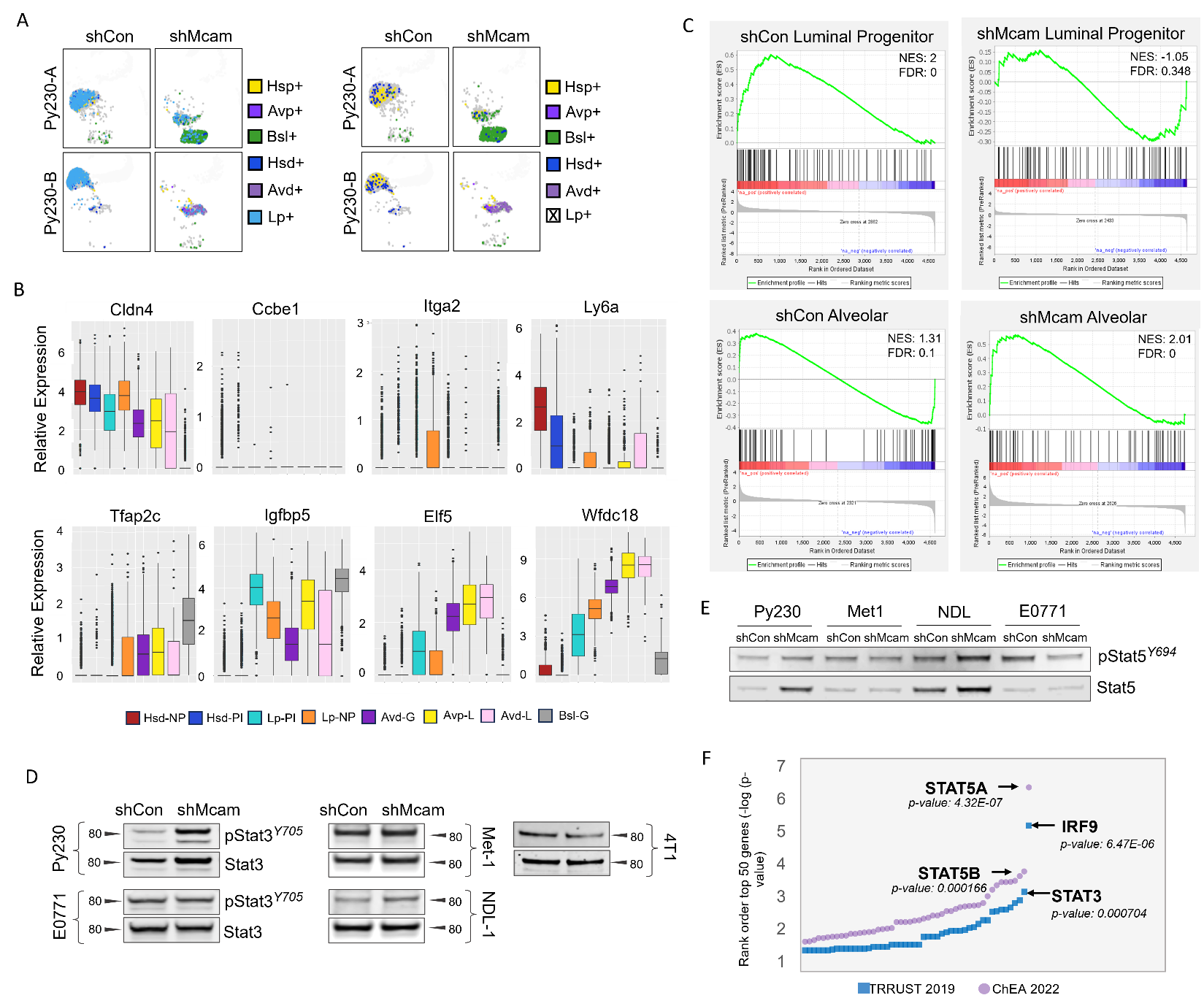
**Supplemental Figure 3.** **Mcam knockdown is associated with increased STAT expression and alveolar phenotypes.** A) Identification of different cell states in Py230 control and Mcam KD cells in clones A and B utilizing gene signatures from (38). B) Select clusters across genes of interest for luminal and alveolar markers utilizing the tool and associated data from [GSE106273](https://www.ncbi.nlm.nih.gov/geo/query/acc.cgi?acc=GSE106273). Hsd=hormone sensing differentiated, Lp=luminal progenitor, Avd=alveolar differentiated, Avp=alveolar progenitor, Bsl=basal, NP=nulliparous, PI=post involution, G=gestation, L=lactation. C) GSEA analysis of Py230 Con and Mcam KD against murine luminal progenitor and alveolar. D) Western blot analysis of pStat3 and total Stat3 across murine mammary cancer cell lines. E) Western blot analysis of pStat5 and total Stat5 across murine mammary cancer cell lines. F) Graphical representation of ChEA 2022 and TRRUST 2019 enrichment data in Py230 Mcam KD cells.

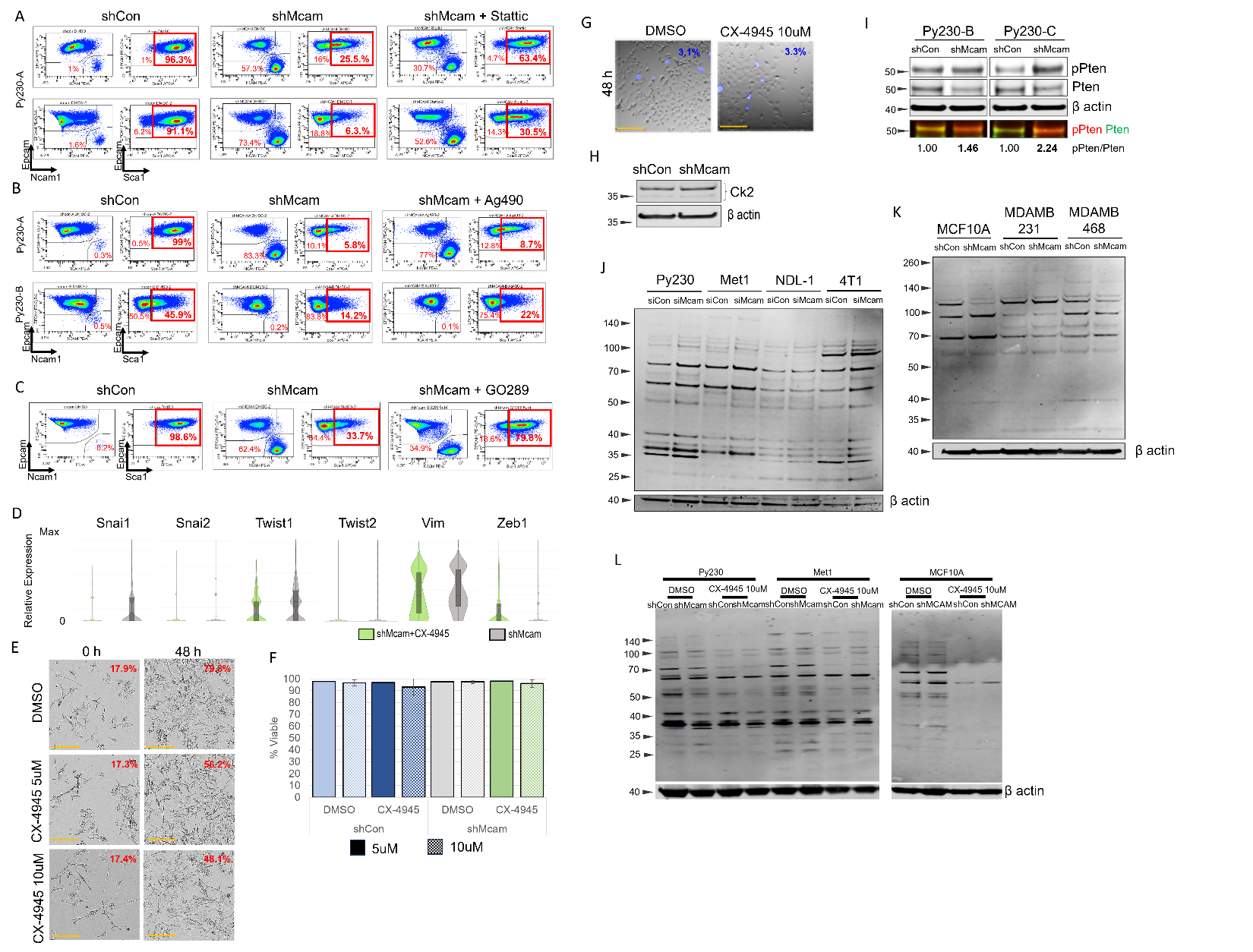
 **Supplemental Figure 4. Stat3 pathway inhibition reverses Mcam KD dependent phenotypes.** A) FACS analysis images of Py230-A cells treated with Stattic 1μM for 7 Days and stained for Epcam, Sca1 and Ncam1. B) FACS analysis images of Py230-A and Py230-B cells treated with 30μM Ag490 for 7 Days and stained for Epcam, Sca1 and Ncam1. C) FACS analysis images of Py230-A cells treated with GO289 5μM for 7 Days and stained for Epcam, Sca1 and Ncam1. D) Py230-A cells treated with CX-4945 10μM for 7 Days and analyzed with scRNA-seq for EMT markers. E) Phase contrast imaging of Py230 parental cells at 0 hour and after 48 hours of treatment with either DMSO vehicle control, 5μM CX-4945 or 10μM CX-4945. Scale bar=100μM. Percentages in red are relative confluency of the culture by Incucyte proliferation analysis. F) Quantification of live cells from E. G) Py230 parental cells after 48 hours of vehicle DMSO treatment or treatment with 10μM CX-4945. Blue percentage is indicative of DAPI+ cells. H) Ck2 protein levels in Py230-A shCon and Mcam KD cells. I) Increased pPten levels in independent clones of Mcam KD Py230. J) Full blot from Figure 4I across multiple murine breast cancer cell lines with transfected siCon or siMcam-A and probed for phosphorylated CK2 substrate protein expression. K) Full blot from Figure 4I across multiple human breast cancer cell lines stably transduced with shCon or shMCAM-A and probed for phosphorylated CK2 substrate protein expression. L) Western blot analysis in select stably transduced shCon/shMCAM murine and human breast cancer cell lines showing differential levels of phosphorylated CK2 substrates that are reduced upon treatment with 10μM CX-4945 compared to DMSO vehicle control.

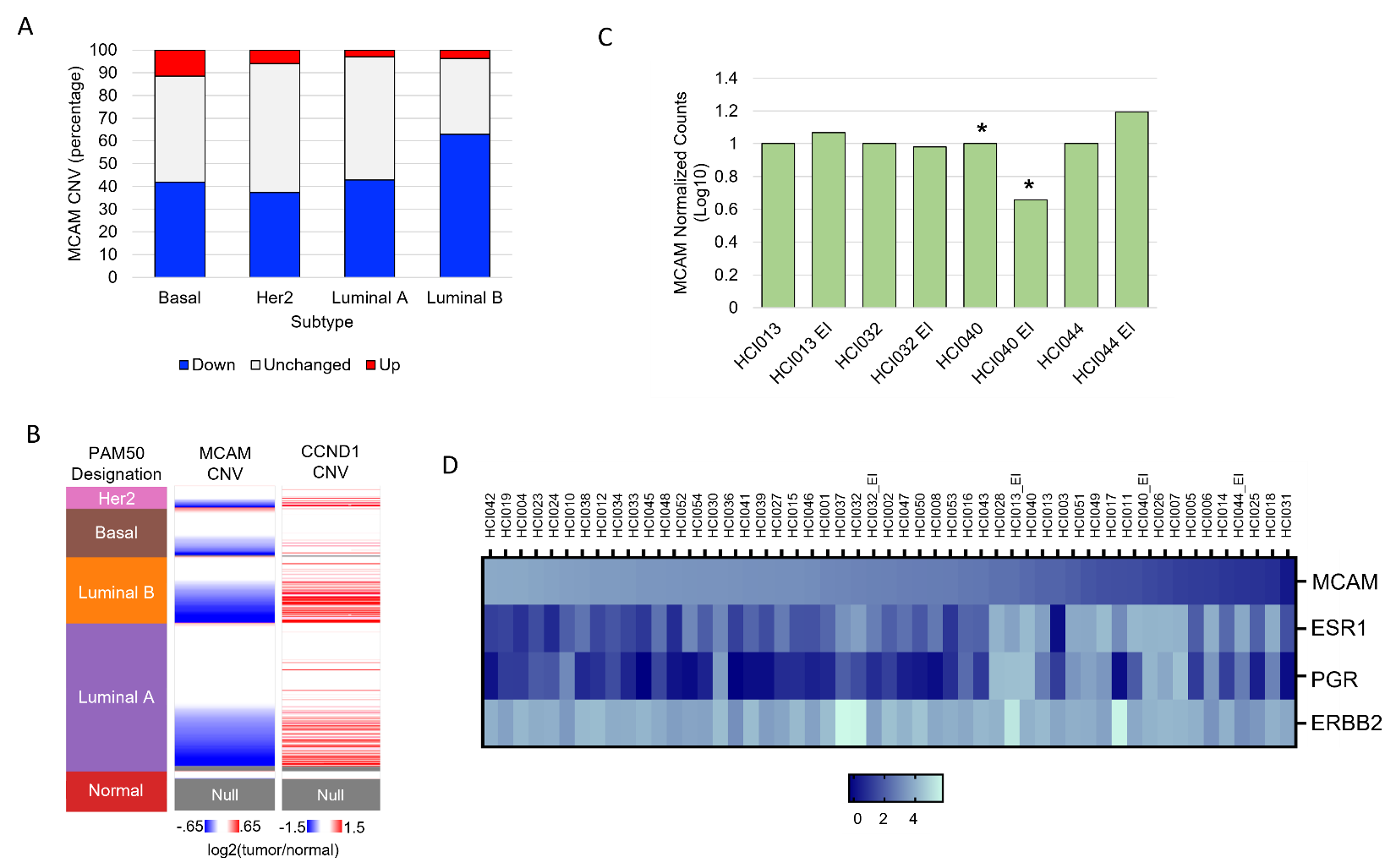

**Supplemental Figure 5. Subtype association of MCAM loss in human breast cancers and patient derived models.** A) TCGA breast cancer data stratified by subtype for MCAM CNV gain or loss. B) TCGA breast cancer data organized by PAM50 designation, expression of PAM50 gene set, MCAM copy number variation, and CCND1 copy number variation. C) Normalized MCAM expression across PDMs and their estrogen insensitive (EI) counterparts. Asterisks indicate the PDM line with 11q deletion. D) Heat map across available PDMs for the log(normalized count) for MCAM, ESR1, PGR, and ERBB2.
